# Predicting Protein-Protein Interactions Using Vectorized Sequence Descriptors and Machine Learning Classifiers

**DOI:** 10.64898/2026.10.21.760860

**Authors:** Ruchika Bhat, Vaishnavi Kumari, Virendra Singh Kaira, Zakeerhussain D Kudari, Sai P. Shankar, G. Jaiganesh

**Author notes:** These authors contributed equally to this work (Ruchika Bhat, Vaishnavi Kumari).

## Abstract

Accurate prediction of protein-protein interactions (PPIs) is fundamental for elucidating cellular mechanisms and accelerating drug discovery. Although experimental techniques provide extensive interaction maps, their high cost and time requirements necessitate robust computational alternatives. In this study, we present an interpretable machine-learning framework that integrates sequence-derived descriptors,specifically 120 Auto-Covariance (AC) features, 686 Conjoint Triad (CT) features, and 1,820 Local Descriptors (LD) features, with classical classification algorithms, including Support Vector Machines (SVM), Random Forests (RF), and XGBoost (XGB). Evaluated on gold-standard sequence-based PPI benchmark dataset using stratified five fold cross-validation, the models achieved robust classification accuracies: 86.92% (SVM), 97.75% (RF), and a peak cross-validation accuracy of 97.97% with XGBoost (Rec: 98.09%, Pr: 98.10%, F1: 98.09%). Furthermore, rigorous evaluation on an independent unseen (Human-PPI) test set demonstrated exceptional generalization capabilities. The Random Forest model emerged as the top-performing architecture, yielding an outstanding accuracy of 99.51%, a precision of 99.72%, a recall of 99.29%, an F1-score of 99.50%, an Area Under the Receiver Operating Characteristic curve (AUROC) of 0.9990, and an Area Under the Precision-Recall Curve (AUPRC) of 0.9989 (closely followed by XGBoost with 98.54% accuracy, AUROC 0.9979, and AUPRC 0.9974). Feature-importance analysis revealed that short-range physicochemical correlations (such as polarity and hydrophobicity lags in Protein 2), regional amino-acid compositions, and specific triad patterns (e.g., A/G/V-M/S/T/Y-M/S/T/Y) drive accurate PPI discrimination. These findings demonstrate the effectiveness of domain-informed, handcrafted sequence features combined with machine-learning classifiers for PPI prediction, providing biologically interpretable insights with minimal computational overhead and enabling efficient, scalable interactome screening.

## 1 Introduction

Proteins are large, essential organic molecules in all living organisms composed of twenty standard amino acids linked together by peptide bonds, forming long chains that fold into specific three-dimensional structures. The unique sequence of amino acids and the resulting structure determine a protein’s function^1^. They are essential for nearly all cellular functions, including metabolic cycles, DNA transcription and replication, and signaling cascades^2^. Proteins rarely function in isolation; rather, they engage with each other to create complex networks, disruption or dysregulation of which is commonly correlated with the onset and development of disease. Therefore, understanding their interactions is crucial for deciphering cellular processes, disease mechanisms, and drug development^3–5^. Protein-protein interaction (PPI) prediction has gained significant importance in molecular biology and bioinformatics. These interactions form complex networks that coordinate everything from cellular communication and immune responses to DNA replication and metabolism^1^. Proteins being the most common drug target, they have immense importance in drug discovery. To identify if a particular protein is a target or not, one has to know the protein hotspots and functional relevance. Protein-protein interactions play a vital role in identifying both these aspects of drug discovery.

Advancements in high-throughput experimental techniques, such as yeast two-hybrid (Y2H) screens, have generated vast amounts of protein-interaction data. Despite their widespread use, these methods exhibit significant limitations. Y2H assays, for example, are frequently time-consuming and labor-intensive, rendering them suboptimal for large-scale or real-time studies. They also generate high rates of false positives and false negatives, largely because artificial in vitro settings fail to recapitulate the native physiological environment of living cells. Additionally, these approaches commonly fail to detect transient, weak, or context-dependent interactions, generating incomplete interactome maps with limited biological context. On the other hand, computational approaches offer a complementary strategy to efficiently predict PPIs and guide experimental validation^6^. Recent advancements in deep learning have significantly transformed protein-protein interaction (PPI) analysis, with GNNs, CNNs, and attention mechanisms applied to prediction, site identification, and network analysis. While these methods reveal complex biological patterns, challenges remain in data quality, integration, and model interpretability^7^.

A few techniques that use the amino acid sequence to directly extract information have recently attracted a lot of attention^6,8–12^. One of the best works among them is an SVM-based approach^13^ created by Shen et al.^8^. In the study, the 20 amino acids were clustered into several classes according to their dipoles and volumes of the side chains. The conjoint triad method abstracts the features of protein pairs based on the classification of amino acids and achieves a good performance with an accuracy of 83.90%. It is most likely caused by the various feature representation techniques and data sources. A technique based on SVM and auto covariance was created by Guo et al. in the other study to extract information about interactions in the sequence’s discontinuous amino acid segments^9^. When used to predict *Saccharomyces cerevisiae* PPIs, their method produced an 86.55% prediction accuracy. In one of the research by You et al.^11^ introduced a hierarchical PCA-EELM model for sequence-based PPI prediction, addressing a key challenge in computational biology. It outperforms SVM-based methods in accuracy and efficiency. Key innovations include multiple sequence-based feature extraction techniques (Autocovariance, Conjoint Triad, Local Descriptor, Moran AutoCorrelation), Principal Component Analysis for dimensionality reduction and ensemble ELM classifiers for improved stability, making it a significant advancement in PPI prediction.

The study introduced, by You et al.^14^, a Multi-scale Local Descriptor (MLD) feature representation for protein sequences, capturing multi-scale local information. Combined with a Random Forest classifier^15^, the method achieves 94.72% accuracy on *Saccharomyces cerevisiae* PPI data and outperforms state-of-the-art methods on the *H. pylori* dataset. To increase speed and performance, the state-of-the-art ensemble learning technique known as eXtreme gradient boosting, or XGBoost^16^, uses gradient boosted decision trees for better speed and performance. By controlling overfitting through a more regularized model formalization, XGBoost outperforms other boosting algorithms. In the related problem of protein-protein interface hotspot prediction, Liu et al.^17^ investigated the use of AdaBoost. In order to construct protein-protein interaction networks for *Leishmania* species using protein structural information data, Vasconcelos et al. employed gradient boosting^18^. For the protein-protein interface hotspot prediction problem, Wang et al. combined XGBoost with a novel PREDHS2 technique^19^ and achieved 81% accuracy.

Even with the achievements of conventional machine learning and deep learning methods, these techniques still have their limits in generalizing to varied data sets, robustness against class imbalance, and reliance on clean structural or experimental data. Deep learning models, in many instances, also suffer from lack of interpretability, and hence the difficulty of being able to recover biologically useful information from the predictions they provide.

To overcome these shortcomings, our research aims to construct a sequence-based machine learning paradigm that is both accurate, interpretable, and generalizable. Rather than depending on intricate and frequently impenetrable deep learning models, we utilize three popular and interpretable classifiers which are Support Vector Machine (SVM), Random Forest (RF), and Extreme Gradient Boosting (XGBoost). These models have been selected because of their successful application to high-dimensional biological datasets, their resilience to overfitting, and their capacity for dealing with nonlinear patterns without demanding structural knowledge. Our strategy is based on utilizing biologically inspired feature engineering approaches, like Auto-Covariance (AC), Conjoint Triad (CT), and Local Descriptors (LD), to acquire meaningful information about protein sequences and represent them as vectorized structures that are tuned for classification. In contrast to most previous work that either does not have comprehensive performance testing on unseen data or is heavily dependent on black-box models, this paper focuses on interpretability and practical use particularly in scenarios where structural data or large-scale labeled data are unavailable or scarce.

Additionally, this study addresses challenges in PPI prediction, such as handling imbalanced datasets, incorporating domain knowledge, and interpreting model predictions. The developed methods were evaluated on benchmark datasets and compared with the existing state-of-the-art approaches.

The outcomes of this study have the potential to contribute to the understanding of protein interaction networks, aid in the discovery of novel drug targets, and provide insights into the molecular basis of diseases^7^. Furthermore, the predictive models developed in this study may serve as valuable tools for researchers in the fields of systems biology, drug discovery, and personalized medicine^7^.

## 2 Materials and Methodology

The proposed methodology begins with a curated protein-protein interaction (PPI) dataset, which is divided into a training and validation set and an independent unseen test set. The training and validation data are evaluated using stratified 5-fold cross-validation to obtain reliable estimates of model performance, while the unseen test set is kept separate for assessing generalization. Three complementary sequence-based descriptors, namely Auto-Covariance (AC), Conjoint Triad (CT), and Local Descriptor (LD), are extracted to represent different physicochemical and sequential characteristics of the proteins.

The extracted features are subsequently used to develop three machine-learning classifiers: Support Vector Machine (SVM), Random Forest (RF), and XGBoost. Their performance is assessed using accuracy, recall, precision, F1-score, AUROC, and AUPRC. Following comparative evaluation on the unseen dataset, the best-performing model is selected as the final predictor and used to classify protein pairs as interacting or non-interacting, providing the final PPI prediction output as depicted in Figure 1. A detailed description of the individual steps involved in the proposed methodology is presented in the following subsections.

**Figure 1.**
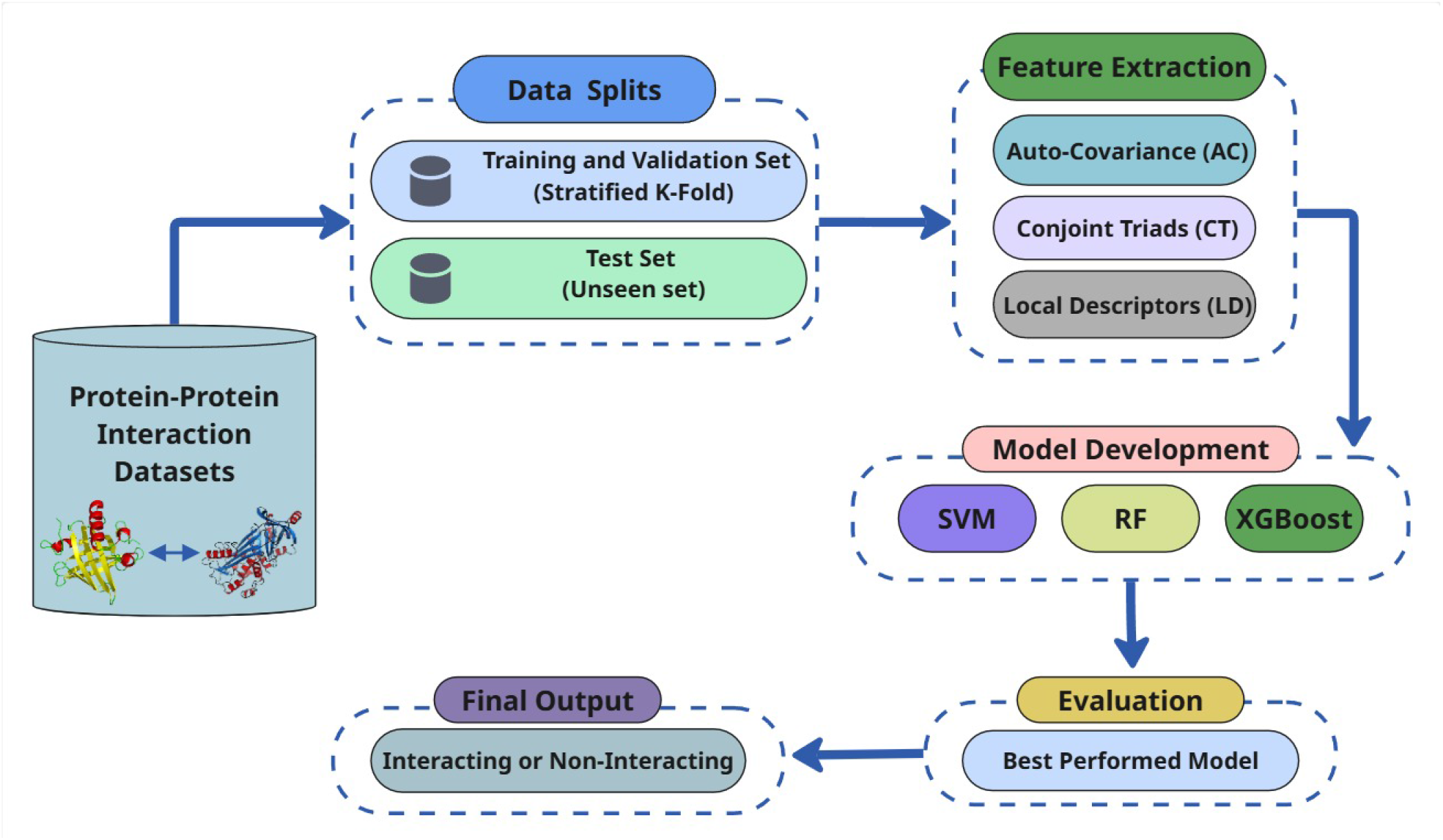
Schematic representation of the proposed machine-learning framework for protein–protein interaction prediction. The workflow begins with dataset splitting and feature extraction using Auto-Covariance, Conjoint Triad, and Local Descriptor methods. These extracted features are used to develop three machine-learning models based on Support Vector Machine (SVM), Random Forest (RF), and XGBoost. The developed models are then evaluated to identify the best-performing model, which is ultimately used to classify protein pairs as interacting or non-interacting.

### 2.1 Dataset

The protein-protein interaction (PPI) dataset used in this study was obtained from the gold-standard sequence-based PPI benchmark developed by Bernett et al.^20^. The dataset is publicly available through Figshare at https://doi.org/10.6084/m9.figshare.21591618. It was specifically developed for sequence-based PPI prediction using experimentally supported human protein interactions and corresponding protein sequences obtained from Swiss-Prot. Importantly, the dataset construction was designed to minimize information leakage between the different subsets by ensuring that proteins assigned to one subset were not present in the other subsets. In addition, KaHIP-based partitioning was employed to minimize sequence similarity between the training, validation, and test partitions, while CD-HIT was used to remove redundant proteins with more than 40% pairwise sequence similarity.

For the present study, the *Intra-0* subset from gold-standard sequence-based PPI was selected because it contains human PPI data and provides a well-defined set of protein pairs for evaluating sequence-based interaction prediction. The subset contains both interacting and non-interacting protein pairs, enabling the formulation of PPI prediction as a binary classification problem.

In addition to the standard training and testing partitions, an independent large-scale prediction of human protein-protein interactions dataset^21^ (Human-PPI) consisting of completely unseen protein pairs was reserved for evaluating model generalizability. These protein pairs were not used during model training or model development, thereby providing a more stringent assessment of the ability of the trained model to generalize to previously unseen interactions. The statistical distribution of the datasets used in this study is summarized in Table 1.

**Table 1.** Statistics of the datasets used for training, testing, and unseen evaluation.

| Type | Dataset Name | Negative | Positive | Total |
| --- | --- | --- | --- | --- |
| Training and Testing Dataset | Gold-standard sequence-based PPI (Intra-0) | 29630 | 29630 | 59260 |
| Unseen Test Dataset | Human-PPI | 36480 | 36480 | 72960 |

### 2.2 Preprocessing

A systematic preprocessing pipeline was applied to the protein-protein interaction (PPI) dataset to ensure data quality, minimize redundancy, prevent information leakage, and generate biologically meaningful numerical representations for machine learning. The preprocessing procedure consisted of the following steps:

1. **Protein Sequence Retrieval and Quality Filtering**. Protein sequences corresponding to the interacting protein pairs were retrieved from the provided Swiss-Prot sequence resource. To remove potentially incomplete or biologically unreliable entries, protein pairs containing sequences shorter than 50 amino acids were excluded from the dataset. This filtering step reduced the influence of fragmented protein sequences and ensured that the subsequent sequence-based feature extraction was performed on sufficiently informative protein sequences.
2. **Removal of Redundant Interactions**. Duplicate interaction records were removed to avoid repeated observations in the dataset. Since PPI relationships are generally undirected, reverse-order duplicates were also identified and eliminated. Thus, interactions represented as A-B and B-A were treated as the same interaction and only a single instance was retained. This step was particularly important for preventing redundancy and potential data leakage between the training and testing datasets.
3. **Dataset Shuffling and Partitioning**. Following quality control and redundancy removal, the curated dataset was randomly shuffled to minimize ordering-related biases. The dataset was subsequently divided using stratified sampling, with 80% of the samples allocated for training and the remaining 20% reserved for testing. Stratification was employed to preserve the original proportion of interacting and non-interacting protein pairs across the two subsets.
4. **Class-Balance Handling**. The curated dataset exhibited an inherent class imbalance, with non-interacting protein pairs being more prevalent than interacting pairs. To reduce the potential bias toward the majority class, a hybrid sampling strategy was employed during model training. This strategy was designed to provide a more balanced representation of interacting and non-interacting pairs while retaining the available information in the original dataset. The statistics of the final preprocessed dataset are presented in Table 2.

**Table 2.**
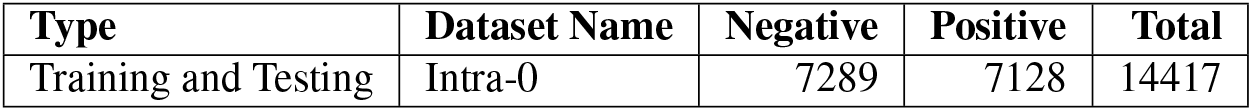
Statistics of the dataset after preprocessing steps.

### 2.3 Feature Extraction

One of the important computational issues in using machine learning for protein-protein interaction (PPI) prediction is to obtain feature vectors that can capture the important biological information hidden in protein sequences. Although a variety of handcrafted descriptors have been put forward, e.g., Amino Acid Composition (AAC), Dipeptide Composition (DPC), Tripeptide Composition (TPC), and Pseudo Amino Acid Composition (PseAAC)^22^, we chose three well-established descriptors: Auto-Covariance (AC), Conjoint Triad (CT), and Local Descriptors (LD). This selection was directed by their differing strengths and overall capacity to incorporate various and biologically informative sequence features, such as global sequence-order dependencies (AC), local physicochemical interactions (CT), and patterns of spatial distributions of amino acids (LD). Collectively, these approaches cover a complete and information-rich set of protein sequence features, ensuring that they are both effective and computationally parsimonious to predicting PPIs.

#### 2.3.1 Auto-Covariance (AC) Scores

The Auto-Covariance (AC) method captures sequence patterns by quantifying the covariation between physicochemical properties of amino acid pairs at varying sequence separations (lags). Our implementation analyzes six key physicochemical descriptors that collectively characterize amino acid interactions across the protein sequence: hydrophobicity (H), polarity (P1), polarizability (P2), side chain volume (VSC), solvent-accessible surface area (SASA), and the net charge index of side chains (NCISC)^23^. These descriptors serve as the foundation for predicting protein-protein interactions. The values of the six physicochemical properties for each amino acid were presented in Table 3^23^. This method was initially employed to convert the amino acid residues into numerical values representing their physicochemical characteristics, subsequently normalized using an equation to achieve a zero mean and a unit standard deviation. This conversion process ensures that the data is standardized, facilitating more accurate comparisons and analyses across different amino acids and their properties.

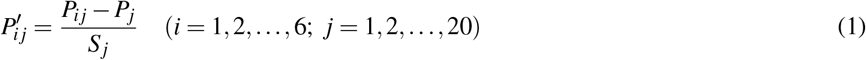

where 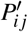 represents the *j*th descriptor value for the *i*th amino acid, *P*_*j*_ denotes the average value of the descriptor calculated across all 20 standard amino acids, and *S* _*j*_ stands for the standard deviation associated with that descriptor. Then each protein sequence was subsequently converted into six separate vectors, where every amino acid was encoded using its normalized property values. These numerical vectors were then processed using the autocovariance (AC) formula to produce fixed-size matrices. For a protein sequence *P* of length *L*, the AC features were computed using the following formula:

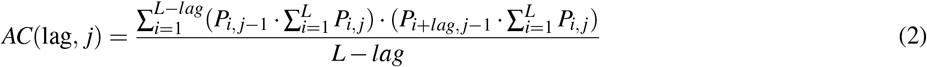

where *lag* refers to the spacing between amino acid residues, *j* indicates the index of one of the physicochemical properties mentioned above, and *i* denotes the position of an amino acid within the sequence *P*.

**Table 3.** Values of the six physicochemical properties of each amino acid.

| Amino Acid | H | VSC | P1 | P2 | SASA | NCISC |
| --- | --- | --- | --- | --- | --- | --- |
| A | 0.62 | 27.5 | 8.1 | 0.046 | 1.181 | 0.007187 |
| C | 0.29 | 44.6 | 5.5 | 0.128 | 1.461 | -0.03661 |
| D | -0.9 | 40 | 13 | 0.105 | 1.587 | -0.02382 |
| E | -0.74 | 62 | 12.3 | 0.151 | 1.862 | 0.006802 |
| F | 1.19 | 115.5 | 5.2 | 0.29 | 2.228 | 0.037552 |
| G | 0.48 | 0 | 9 | 0 | 0.881 | 0.179052 |
| H | -0.4 | 79 | 10.4 | 0.23 | 2.025 | -0.01069 |
| I | 1.38 | 93.5 | 5.2 | 0.186 | 1.81 | 0.021631 |
| K | -1.5 | 100 | 11.3 | 0.219 | 2.258 | 0.017708 |
| L | 1.06 | 93.5 | 4.9 | 0.186 | 1.931 | 0.051672 |
| M | 0.64 | 94.1 | 5.7 | 0.221 | 2.034 | 0.002683 |
| N | -0.78 | 58.7 | 11.6 | 0.134 | 1.655 | 0.005392 |
| P | 0.12 | 41.9 | 8 | 0.131 | 1.468 | 0.239531 |
| Q | -0.85 | 80.7 | 10.5 | 0.18 | 1.932 | 0.049211 |
| R | -2.53 | 105 | 10.5 | 0.291 | 2.56 | 0.043587 |
| S | -0.18 | 29.3 | 9.2 | 0.062 | 1.298 | 0.004627 |
| T | -0.05 | 51.3 | 8.6 | 0.108 | 1.525 | 0.003352 |
| V | 1.08 | 71.5 | 5.9 | 0.14 | 1.645 | 0.057004 |
| W | 0.81 | 145.5 | 5.4 | 0.409 | 2.663 | 0.037977 |
| Y | 0.26 | 117.3 | 6.2 | 0.298 | 2.368 | 0.023599 |

The number of AC variables, *D*, can thus be computed as *D* = *lg* × *q*, where *q* represents the total number of descriptors, and *lg* denotes the maximum lag value, with lag ranging from 1 to *lg*. Once each protein sequence was encoded as a vector of AC features, the vectors corresponding to a pair of proteins were concatenated to form a single feature vector representing the protein pair.

#### 2.3.2 Conjoint Triads (CT) Scores

The Conjoint Triad (CT) method treats every set of three consecutive amino acids as a single unit, capturing both the properties of a central amino acid and those of its neighboring residues^8^. These triads are categorized based on amino acid groupings. By computing the frequency of each triad type, protein-protein interaction (PPI) data can be mapped into a uniform vector space. Prior to employing this approach, the twenty standard amino acids were partitioned into seven groups according to their side chain volumes and dipole characteristics, as depicted in Table 4^11^. This grouping reduces the complexity of the feature space, resulting in 343 possible triads (7 × 7 × 7). For each protein pair, the CT features of the two sequences were combined to produce a 686-dimensional feature vector.

**Table 4.** Division of amino acids based on dipoles and volumes of the side chains.

| No. | Group |
| --- | --- |
| 1 | A, G, V |
| 2 | C |
| 3 | D, E |
| 4 | F, I, L, P |
| 5 | H, N, Q, W |
| 6 | K, R |
| 7 | M, S, T, Y |

#### 2.3.3 Local Descriptors (LD) Scores

The efficacy of the local descriptor, an alignment-independent approach, is primarily contingent on the underlying amino acid groupings. To streamline the representation of the 20 standard amino acids, we initially grouped them into seven functional categories based on their side chain volumes and dipole moments using Table 4. Three local descriptors were subsequently computed based on fluctuations in the occurrence of these amino acid groups within the primary protein sequence: Composition (C), Transition (T), and Distribution (D). C describes the local fractional abundance of each amino acid group. T represents the percentage frequency with which an amino acid from one group is followed by an amino acid from another group. D characterizes the spread of residues in a group by calculating their positions at the 1st, 25th, 50th, 75th, and 100th percentiles within the sequence. If the descriptors are computed over the entire sequence, the resulting feature vector comprises 910 features (7 composition, 49 transition, and 35 distribution).

To effectively capture both continuous and discontinuous protein-protein interaction information, the protein sequence is divided into 10 distinct local regions (A-J), each with unique lengths and structural positions. The protein sequence is divided into four equal-length regions, which are then called regions A, B, C, and D. By splitting the protein sequence into two equal-length sections, regions E and F are produced. Region G, which contains 50% of the sequence, is in the middle. The first 75% of the sequence is represented by Region H, the last 75% by Region I, and the middle 50% by Region J. Figure 2 shows examples of these regions. Each of the 10 segments yields a 91-dimensional local descriptor, generating a 910-dimensional feature vector per protein. For protein-protein interactions, the individual vectors of both proteins are concatenated, producing a final 1820-dimensional combined feature representation.

**Figure 2.**
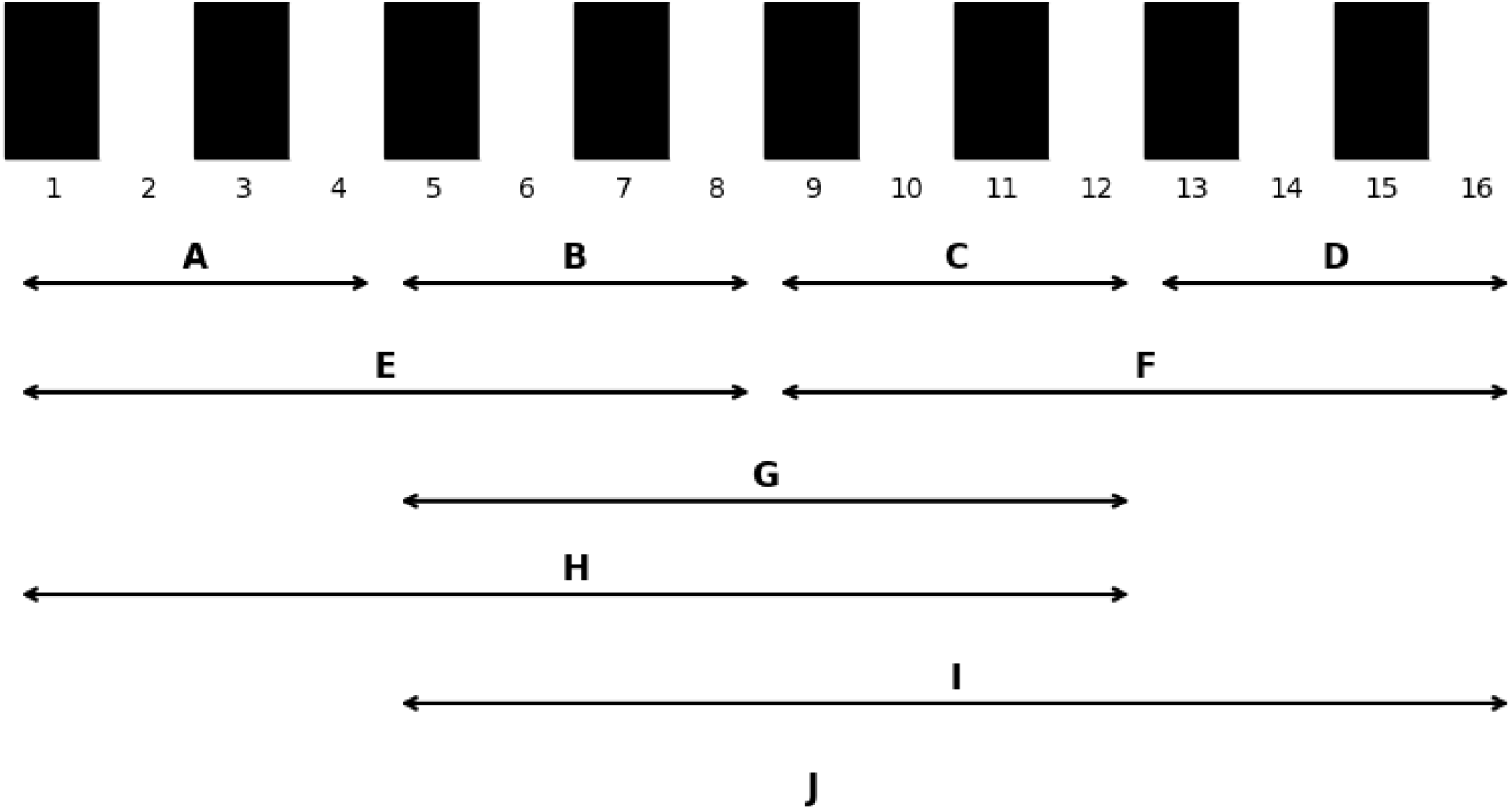
The 10 regions (A-J) used by the local descriptor technique for a theoretical protein sequence. The regions A-D and E-F are obtained by dividing the entire sequence into four equal regions and into two equal regions, I and II, respectively. Region G represents the central 50% of the sequence. Regions H, I, and J are the first, final, and central 75% of the sequence.

#### 2.3.4 Feature Space

The feature set constructed for each protein pair comprised three complementary sequence-derived descriptor groups: auto-covariance (AC), conjoint triad (CT), and local descriptor (LD) features. In total, each sample was represented by 2,626 features, including 120 AC, 686 CT, and 1,820 LD features as shown in Table 5. These descriptors capture complementary aspects of protein sequence characteristics, including sequence-order correlations, local amino acid patterns, and long-range distributional properties. The resulting feature representation was subsequently used as the input for machine-learning-based PPI prediction. Detailed feature ranges and their corresponding mappings are provided in Tables 6, 7, and 8.

**Table 5.**
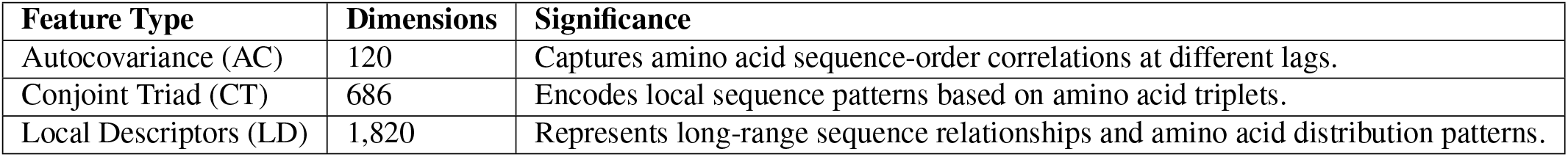
Composition of sequence-derived features used for protein-protein interaction prediction.

**Table 6.** Feature-index ranges and corresponding physicochemical descriptors for the Autocovariance (AC) feature set.

| Feature Range | Property | Protein | Lag Range | Description |
| --- | --- | --- | --- | --- |
| 1-10 | H | Protein 1 | 1-10 | Hydrophobicity autocovariance |
| 11-20 | H | Protein 2 | 1-10 | Hydrophobicity autocovariance |
| 21-30 | VSC | Protein 1 | 1-10 | Van der Waals steric coefficient autocovariance |
| 31-40 | VSC | Protein 2 | 1-10 | Van der Waals steric coefficient autocovariance |
| 41-50 | P1 | Protein 1 | 1-10 | Polarity autocovariance |
| 51-60 | P1 | Protein 2 | 1-10 | Polarity autocovariance |
| 61-70 | P2 | Protein 1 | 1-10 | Polarity autocovariance |
| 71-80 | P2 | Protein 2 | 1-10 | Polarity autocovariance |
| 81-90 | SASA | Protein 1 | 1-10 | Solvent-accessible surface area autocovariance |
| 91-100 | SASA | Protein 2 | 1-10 | Solvent-accessible surface area autocovariance |
| 101-110 | NCISC | Protein 1 | 1-10 | NCISC autocovariance |
| 111-120 | NCISC | Protein 2 | 1-10 | NCISC autocovariance |

**Table 7.** Feature-index ranges and corresponding amino acid group combinations for Conjoint Triad (CT) descriptors. The CT descriptors occupy features 121-806 after the 120 Autocovariance (AC) features.

| Feature Range | Protein | Group 1 | Group 2 | Group 3 |
| --- | --- | --- | --- | --- |
| 121-127 | Protein 1 | 1 (A,G,V) | 1 (A,G,V) | 1-7 |
| 128-134 | Protein 1 | 1 (A,G,V) | 2 (C) | 1-7 |
| 135-141 | Protein 1 | 1 (A,G,V) | 3 (D,E) | 1-7 |
| 142-148 | Protein 1 | 1 (A,G,V) | 4 (F,I,L,P) | 1-7 |
| 149-155 | Protein 1 | 1 (A,G,V) | 5 (H,N,Q,W) | 1-7 |
| 156-162 | Protein 1 | 1 (A,G,V) | 6 (K,R) | 1-7 |
| 163-169 | Protein 1 | 1 (A,G,V) | 7 (M,S,T,Y) | 1-7 |
| 170-218 | Protein 1 | 2 (C) | 1-7 | 1-7 |
| 219-267 | Protein 1 | 3 (D,E) | 1-7 | 1-7 |
| 268-316 | Protein 1 | 4 (F,I,L,P) | 1-7 | 1-7 |
| 317-365 | Protein 1 | 5 (H,N,Q,W) | 1-7 | 1-7 |
| 366-414 | Protein 1 | 6 (K,R) | 1-7 | 1-7 |
| 415-463 | Protein 1 | 7 (M,S,T,Y) | 1-7 | 1-7 |
| 464-806 | Protein 2 | Same combinations | Same combinations | 1-7 |

**Table 8.** Feature-index ranges and sequence regions represented by the Local Descriptor (LD) features. The LD descriptors occupy features 807-2626 following the AC and CT descriptors.

| Feature Range | Protein | Region | Sequence Coverage |
| --- | --- | --- | --- |
| 807-897 | Protein 1 | A | First quarter (0-25%) |
| 898-988 | Protein 1 | B | Second quarter (25-50%) |
| 989-1079 | Protein 1 | C | Third quarter (50-75%) |
| 1080-1170 | Protein 1 | D | Last quarter (75-100%) |
| 1171-1261 | Protein 1 | E | First half (0-50%) |
| 1262-1352 | Protein 1 | F | Last half (50-100%) |
| 1353-1443 | Protein 1 | G | Middle 50% (25-75%) |
| 1444-1534 | Protein 1 | H | First 75% (0-75%) |
| 1535-1625 | Protein 1 | I | Last 75% (25-100%) |
| 1626-1716 | Protein 1 | J | Middle 75% (12.5-87.5%) |
| 1717-1807 | Protein 2 | A | First quarter (0-25%) |
| 1808-1898 | Protein 2 | B | Second quarter (25-50%) |
| 1899-1989 | Protein 2 | C | Third quarter (50-75%) |
| 1990-2080 | Protein 2 | D | Last quarter (75-100%) |
| 2081-2171 | Protein 2 | E | First half (0-50%) |
| 2172-2262 | Protein 2 | F | Last half (50-100%) |
| 2263-2353 | Protein 2 | G | Middle 50% (25-75%) |
| 2354-2444 | Protein 2 | H | First 75% (0-75%) |
| 2445-2535 | Protein 2 | I | Last 75% (25-100%) |
| 2536-2626 | Protein 2 | J | Middle 75% (12.5-87.5%) |

**Table 9.** Composition of the 91 Local Descriptor features calculated for each sequence region.

| Component | Features | Range within Region | Description |
| --- | --- | --- | --- |
| Composition | 7 | 1-7 | Frequency of the seven amino acid groups |
| Transition | 49 | 8-56 | Pairwise transitions between the seven amino acid groups |
| Distribution | 35 | 57-91 | Five positional distribution measures for each group |
| **Total** | **91** | **1-91** | Complete descriptor for one sequence region |

### 2.4 Algorithms Used for PPI Prediction

Several classification models were trained using the extracted features to predict PPI. The following machine learning algorithms were employed: SVM, RF, and XGB, and to improve model generalization, stratified K-fold cross-validation were used during the implementation. The dataset was split into 5 folds, ensuring robustness in model training and evaluation. Using the preprocessed and balanced dataset, each of these algorithms was trained. Below is a detailed explanation of how each algorithm works, along with its mathematical formulation and usage in PPI prediction.

#### 2.4.1 Support Vector Machine (SVM)

A support vector machine model was employed to differentiate between interacting and non-interacting protein pairs. SVM is an ML algorithm utilized for classification, regression, and outlier detection tasks. SVM finds the hyperplane with the greatest separation between classes, optimizing the margin. It excels at dealing with multidimensional data and nonlinear relationships.

Given a dataset {(*x*_*i*_, *y*_*i*_)}_*i*=1,2,3,…,*n*_ where *x*_*i*_ represents feature vectors and *y*_*i*_ is the label (1 for interacting and 0 for non-interacting protein pairs), the SVM model optimizes the following objective:

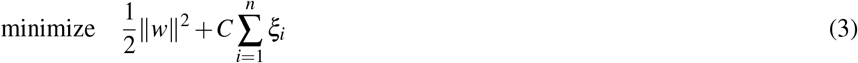

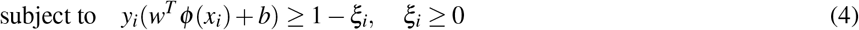

where *w* is the weight vector, *b* is the bias, and *ϕ* (*x*) is the feature transformation function and *ξ*_*i*_ are slack variables allowing soft-margin classification. The trade-off between minimizing classification error and maximizing the margin is controlled by the parameter *C*.

#### 2.4.2 Random Forest (RF)

The Random Forest (RF) model was employed to model complex, non-linear patterns within the protein interaction features. As an ensemble method, it builds multiple decision trees, each trained on randomly selected subsets of the data.

The prediction of an RF model is obtained by averaging the outputs of individual decision trees:

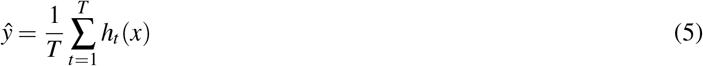

where *h*_*t*_(*x*) denotes the prediction made by the *t*-th decision tree, and *T* is the total number of trees.

#### 2.4.3 XGBoost

XGBoost (eXtreme Gradient Boosting) improves prediction performance by refining the boosting algorithm. In contrast to RF, which builds multiple independent trees, XGB creates trees in sequence, with each new tree aiming to reduce the errors made by the ones before it.

XGBoost minimizes a regularized loss function:

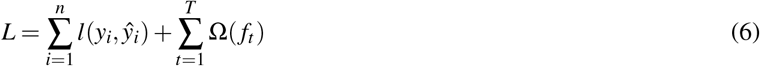

where Ω(*f*_*t*_) is the regularization term governing model complexity and *l*(·) is the loss function.

## 3 Results and Discussion

This section examines how well the proposed PPI prediction techniques perform using a range of assessment metrics. These metrics include Accuracy (Acc), Precision (Pr), Recall (Rc), F1-score, the area under the precision-recall curve (AUPRC), and the area under the receiver operating characteristic curve (AUROC).

### 3.1 Experimental Setup

The proposed protein-protein interaction (PPI) prediction techniques were implemented using Scikit-learn, which provided the machine-learning algorithms and utilities for model development, training, evaluation, and feature-importance analysis. The experiments were conducted on a linux-based machine having NVIDIA RTX 6000 AD GPUs (48GB VRAM) with 128GB of system RAM.

### 3.2 Evaluation Metrics

To evaluate the performance of the proposed protein-protein interaction (PPI) prediction models, two complementary evaluation strategies were employed: stratified five-fold cross-validation and independent hold-out test-set evaluation. These approaches were selected to assess both the robustness of the models during training and their ability to generalize to previously unseen protein pairs.

First, the labeled dataset was divided into training and testing subsets using an 80:20 split. The training portion was further evaluated using stratified five-fold cross-validation, in which the data were divided into five subsets while preserving the proportion of interacting and non-interacting protein pairs in each fold. In each iteration, four folds were used for model training and the remaining fold was used for validation. This process was repeated five times such that each fold was used once for validation. The resulting performance measures were averaged across the five folds to obtain a more reliable estimate of model performance and reduce the effect of variation caused by a particular train-validation partition.

Second, an independent hold-out test set was used for generalization evaluation. The test samples were not used during model training, cross-validation, or model selection. Consequently, performance on this set provides an assessment of how well the trained models can predict previously unseen protein pairs and approximates their expected performance on new data.

Six evaluation metrics were used to quantify the predictive performance of the proposed models: accuracy (Acc), recall (Rec), precision (Pr), F1-score (F1), area under the receiver operating characteristic curve (AUROC), and area under the precision-recall curve (AUPRC). These metrics collectively evaluate overall classification correctness, identification of interacting protein pairs, the reliability of positive predictions, and threshold-independent discrimination performance.

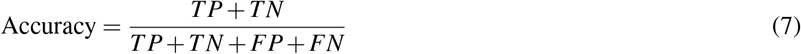

Accuracy measures the proportion of correctly classified protein pairs among all evaluated samples.

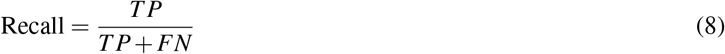

Recall, also referred to as sensitivity or the true positive rate (TPR), measures the proportion of actual interacting protein pairs that are correctly identified by the model.

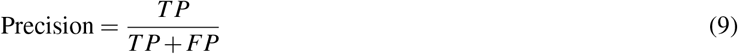

Precision measures the proportion of protein pairs predicted as interacting that are actually positive interactions.

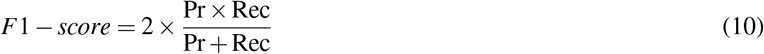

The F1-score represents the harmonic mean of precision and recall and provides a combined measure of the model’s ability to identify positive interactions while limiting false-positive predictions.

The receiver operating characteristic (ROC) curve describes the relationship between the true positive rate (TPR) and the false positive rate (FPR) over different classification thresholds. The area under this curve is calculated as:

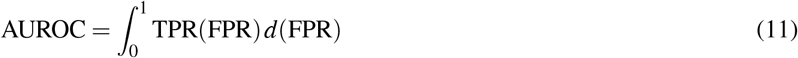

AUROC measures the model’s ability to distinguish between interacting and non-interacting protein pairs across different decision thresholds. A higher AUROC indicates better discrimination between the two classes.

The precision-recall curve represents the relationship between precision and recall across different classification thresholds. Its corresponding area under the curve is given by:

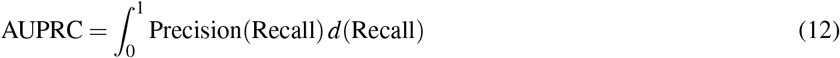

AUPRC summarizes the trade-off between precision and recall and is particularly informative when the distribution of interacting and non-interacting protein pairs is imbalanced.

The false positive rate (FPR) and true positive rate (TPR) used in the ROC analysis are defined as:

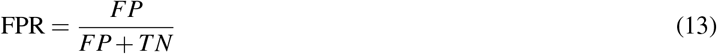

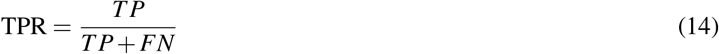

Here, *TP* (true positive) denotes the number of interacting protein pairs correctly predicted as interacting, whereas *TN* (true negative) represents the number of non-interacting protein pairs correctly predicted as non-interacting. *FP* (false positive) denotes the number of non-interacting protein pairs incorrectly predicted as interacting, while *FN* (false negative) represents the number of interacting protein pairs incorrectly predicted as non-interacting. Together, these measures provide a comprehensive assessment of the predictive performance and generalization capability of the proposed PPI prediction models.

### 3.3 Prediction Performance

#### 3.3.1 Cross-Validation Results

The five-fold cross-validation results demonstrate strong predictive performance across all evaluated machine-learning models. SVM achieved an accuracy of 86.92%, with comparable recall, precision, and F1-score of 86.66%, 86.96%, and 86.62%, respectively. RF substantially improved performance, attaining 97.75% accuracy and F1-score, with 97.75% recall and 97.77% precision. XGB yielded the highest overall performance, achieving 97.97% accuracy, 98.09% recall, 98.10% precision, and 98.09% F1-score. These results indicate that ensemble-based approaches, particularly XGB, provide highly consistent and effective predictions compared with SVM. The average performance of each classifier across the 5 folds is summarized in Table 10.

**Table 10.** Average performance metrics over 5-fold cross-validation.

| Models/Metrics | Acc (%) | Rec (%) | Pr (%) | F1-Score (%) |
| --- | --- | --- | --- | --- |
| SVM | 86.92 | 86.66 | 86.96 | 86.62 |
| RF | 97.75 | 97.75 | 97.77 | 97.75 |
| XGB | **97.97** | **98.09** | **98.10** | **98.09** |

#### 3.3.2 Performance on Unseen Data

To evaluate the model’s real-world applicability, each trained model was tested on a completely unseen dataset. SVM achieved 87.58% accuracy, with an F1-score of 87.53%, AUROC of 0.9499, and AUPRC of 0.9484. RF showed consistently high performance, achieving 99.51% accuracy, 99.29% recall, 99.72% precision, and a 99.50% F1-score. It also attained AUROC and AUPRC values of 0.9990 and 0.9989, respectively. XGB similarly performed strongly, with 98.54% accuracy and an F1-score of 98.53%, alongside AUROC and AUPRC values of 0.9979 and 0.9974. Overall, all models demonstrated robust predictive performance on unseen data.

The consistent performance on the holdout test set confirms that the model avoids overfitting and captures generalizable features from the protein sequence data. The results are summarized in Table 11.

**Table 11.** Model performance on unseen data.

| Model/Metrics | Acc (%) | Rec (%) | Pr (%) | F1-Score (%) | AUROC | AUPRC |
| --- | --- | --- | --- | --- | --- | --- |
| SVM | 87.58 | 87.50 | 87.92 | 87.53 | 0.9499 | 0.9484 |
| RF | **99.51** | **99.29** | **99.72** | **99.50** | **0.9990** | **0.9989** |
| XGB | 98.54 | 98.13 | 98.94 | 98.53 | 0.9979 | 0.9974 |

Based on the unseen data evaluation, RF demonstrates the strongest overall performance, achieving the highest accuracy, precision, recall, F1-score, AUROC, and AUPRC. Its consistently superior results indicate robust generalization and reliable classification performance. Therefore, RF was selected as the final model for subsequent protein–protein interaction prediction and downstream analysis.

#### 3.3.3 ROC and Precision-Recall Curve Analysis

To further assess the discriminative performance of the developed classifiers, Receiver Operating Characteristic (ROC) and Precision-Recall (PR) curves were evaluated for Support Vector Machine (SVM), Random Forest (RF), and XGBoost (XGB) across varying decision thresholds. The Area Under the ROC Curve (AUROC) provides an overall measure of the ability to distinguish interacting from non-interacting protein pairs, whereas the Area Under the Precision-Recall Curve (AUPRC) summarizes the trade-off between precision and recall and is particularly informative for evaluating classifier performance under class imbalance. The AUROC and AUPRC values obtained on the independent unseen test set are presented in Table 11.

Among the evaluated classifiers, RF achieved the highest AUROC (0.9990) and AUPRC (0.9989) as shown in Figure 3, demonstrating excellent discrimination between interacting and non-interacting PPI pairs and a strong balance between precision and recall across classification thresholds.

**Figure 3.**
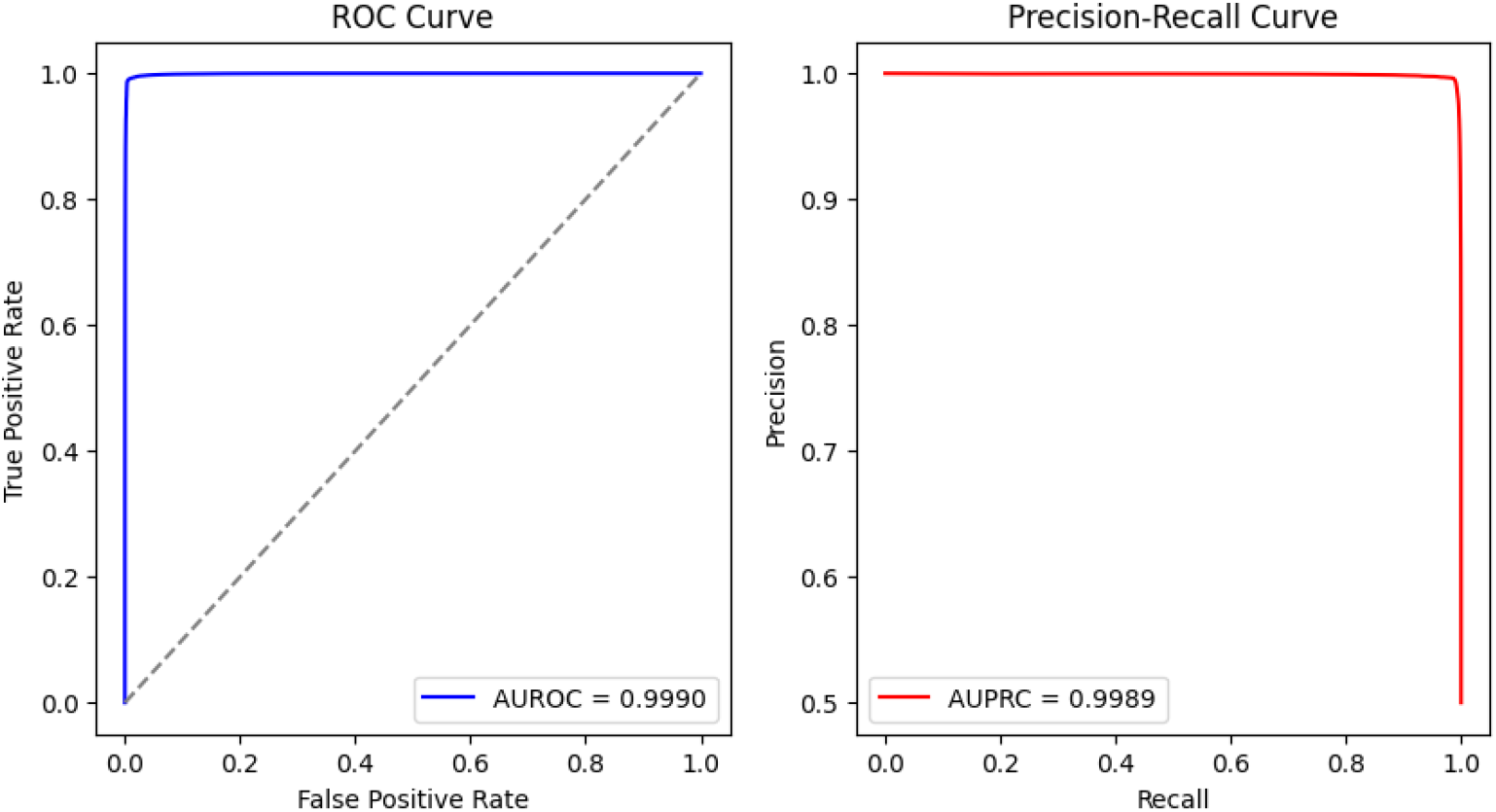
Receiver Operating Characteristic (ROC) and Precision-Recall (PR) curves illustrating the performance of the Random Forest model on the independent unseen test set (AUROC = 0.9990, AUPRC = 0.9989).

XGB also exhibited highly competitive performance, with AUROC and AUPRC values of 0.9979 and 0.9974, respectively as depicted in Figure 4, indicating strong discriminative capability on unseen protein pairs.

**Figure 4.**
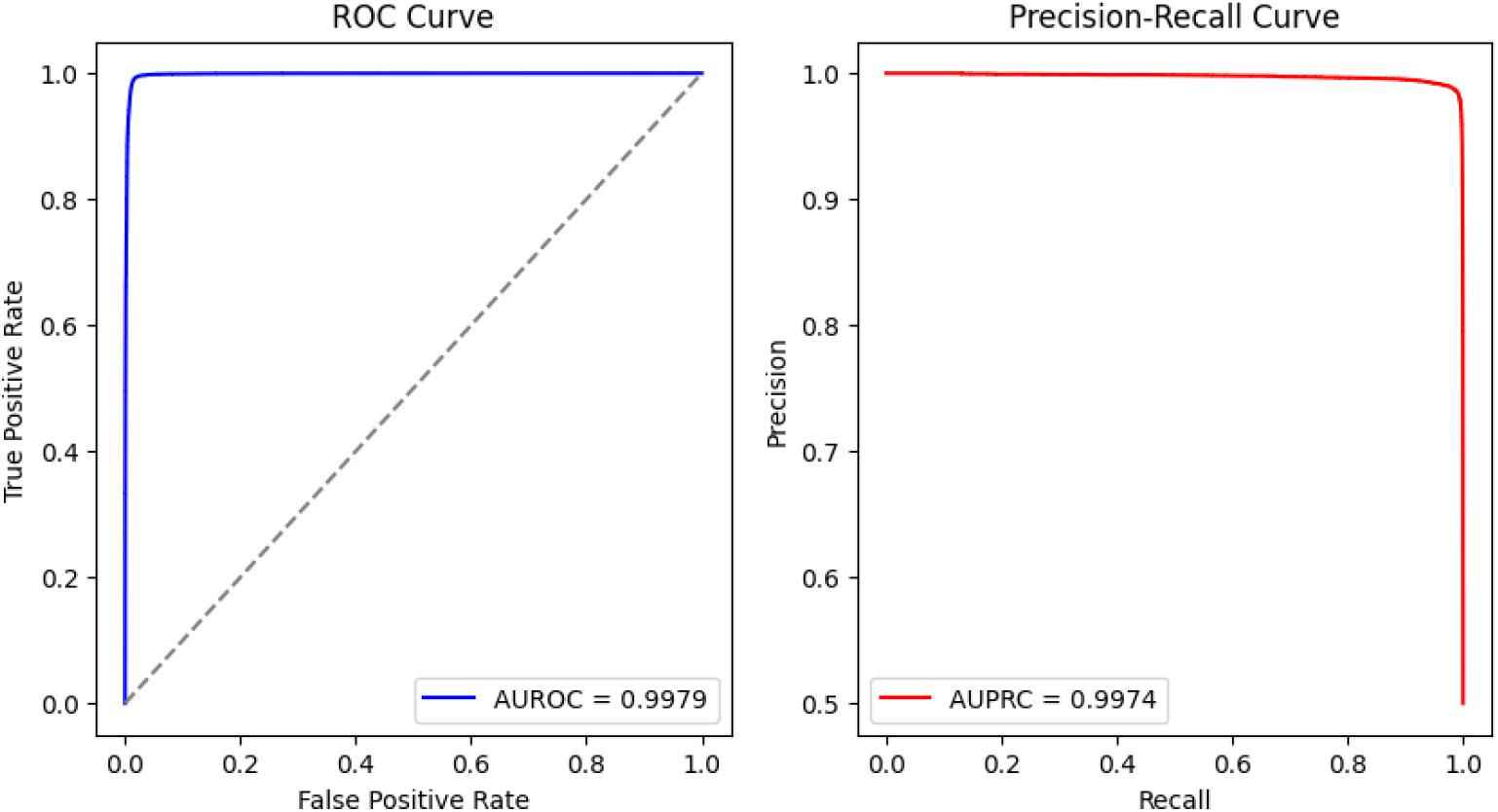
Receiver Operating Characteristic (ROC) and Precision-Recall (PR) curves illustrating the performance of the XGBoost model on the independent unseen test set (AUROC = 0.9979, AUPRC = 0.9974).

In comparison, SVM achieved AUROC and AUPRC values of 0.9499 and 0.9484, respectively. Although SVM demonstrated strong predictive performance can be interpreted from Figure 5, its comparatively lower scores suggest that it may be less effective than the tree-based ensemble approaches in capturing complex nonlinear relationships within the high-dimensional feature space. The consistently high AUROC and AUPRC values of RF and XGB indicate ensemble based models ability to maintain strong classification performance across different decision thresholds. Overall, the ROC and PR analysis further support the robust generalization of the evaluated models, with RF achieving the highest performance on the independent unseen dataset.

**Figure 5.**
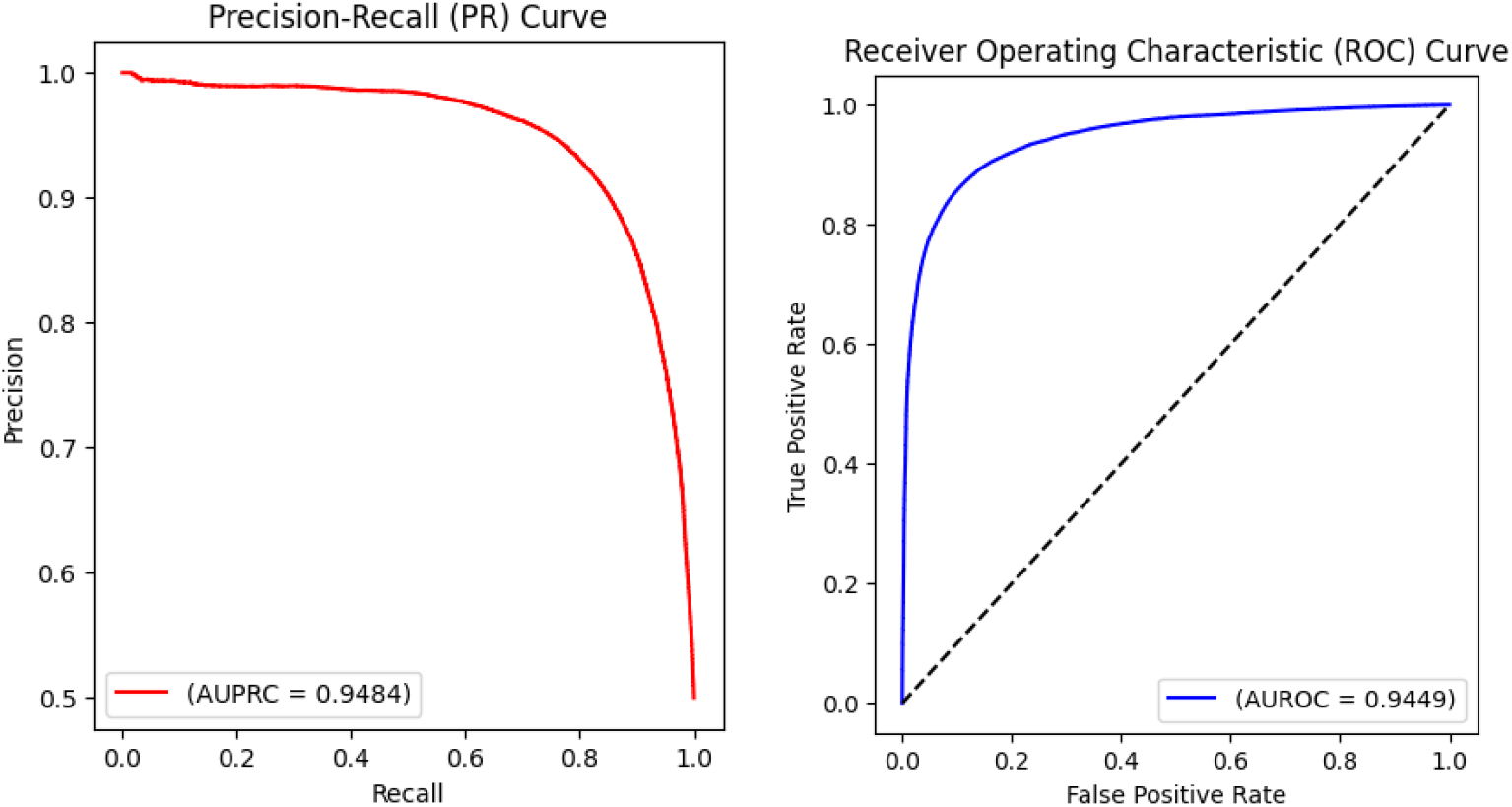
Receiver Operating Characteristic (ROC) and Precision-Recall (PR) curves illustrating the performance of the SVM model on the independent unseen test set.

#### 3.3.4 Interpretability and feature importance

Feature-importance analysis improves the interpretability of machine learning models by identifying the input features that contribute most strongly to model predictions. In PPI prediction, it helps to reveal which protein sequence descriptors are most informative, providing insights into potentially relevant biological patterns. This analysis bridges model performance and biological interpretation, making predictions more transparent, explainable, and easier to validate. Detailed feature ranges and their corresponding mappings are provided in Tables 6, 7, and 8.

Random Forest feature-importance analysis identified a combination of physicochemical auto-covariance (AC), conjoint triad (CT), and local composition-transition-distribution (LD) descriptors among the most informative features for PPI prediction, as shown in Figure 6 and summarized in Table 12. The five highest-ranked features were indices 52, 12, 512, 2086, and 16. Feature 52, the most important descriptor, corresponded to the polarity (P1) autocovariance of Protein 2 at a sequence lag of two, indicating that short-range polarity organization may contribute substantially to PPI discrimination. Feature 12 represented the hydrophobicity (H) autocovariance of Protein 2 at lag two, further highlighting the importance of short-range physicochemical correlations. The third-ranked feature, index 512, was a conjoint triad descriptor corresponding to the A/G/V-M/S/T/Y-M/S/T/Y residue-group pattern in Protein 2, suggesting that specific local three-residue sequence patterns also contribute to the model’s predictions. Feature 2086 represented the composition of K/R residues in the first 50% of Protein 2, demonstrating the contribution of regional amino-acid composition. Finally, feature 16 corresponded to hydrophobicity autocovariance of Protein 2 at lag six. Collectively, the top five features indicate that the Random Forest model primarily exploits complementary information from short-range physicochemical correlations, local three-residue patterns, and region-specific amino-acid composition to distinguish interacting from non-interacting protein pairs.

**Table 12.** Top five Random Forest features identified by feature-importance analysis.

| Rank | Feature Index | Type | Feature Description |
| --- | --- | --- | --- |
| 1 | 52 | AC | P1 autocovariance of Protein 2 at lag 2 |
| 2 | 12 | AC | H autocovariance of Protein 2 at lag 2 |
| 3 | 512 | CT | A/G/V-M/S/T/Y-M/S/T/Y triad pattern in Protein 2 |
| 4 | 2086 | LD | K/R composition in the first 50% of Protein 2 |
| 5 | 16 | AC | H autocovariance of Protein 2 at lag 6 |

**Figure 6.**
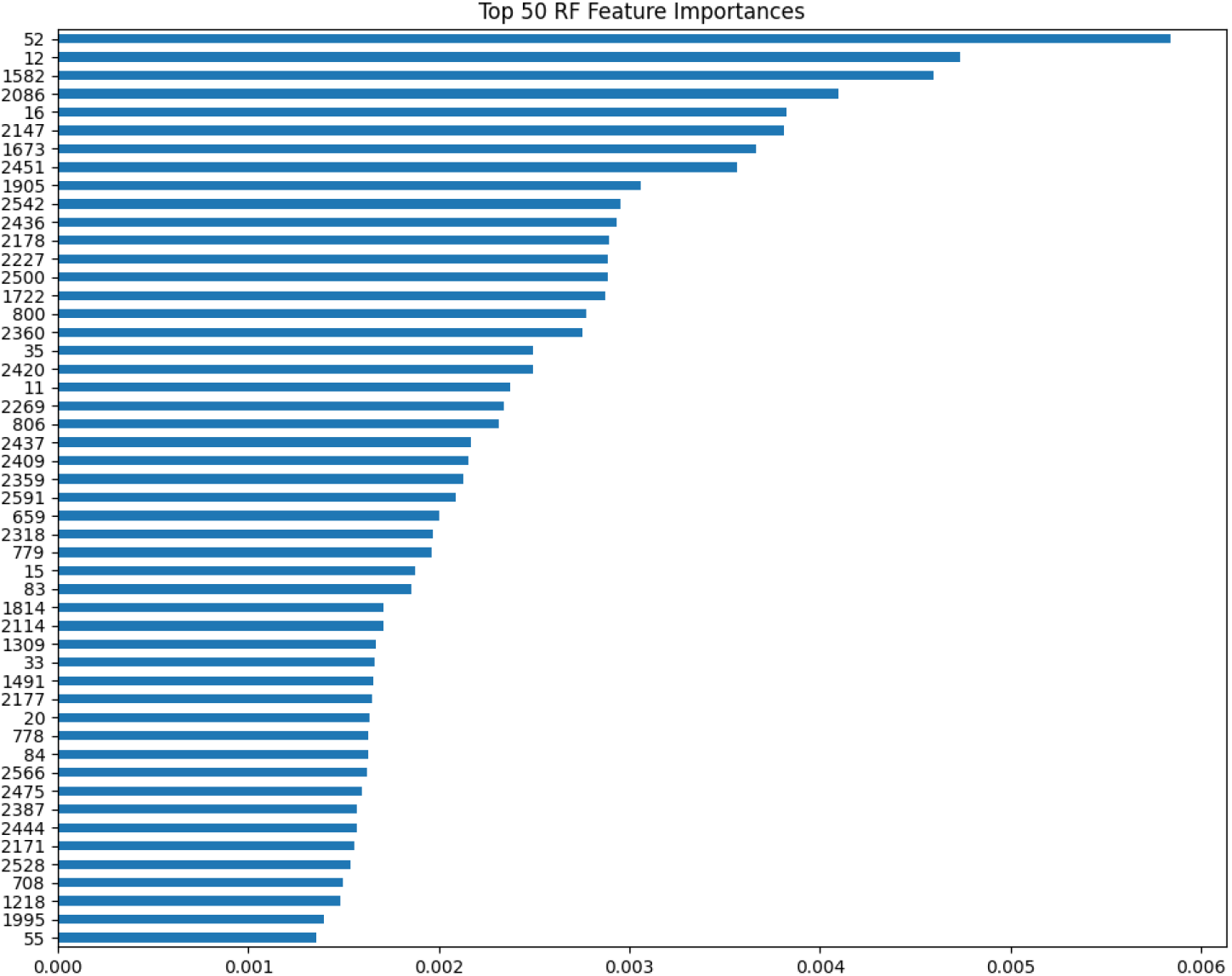
Top 50 Random Forest features ranked by mean feature importance, highlighting key Autocovariance (AC), Conjoint triad (CT), and Local Descriptors (LD) contributing to PPI prediction.

The XGBoost feature-importance analysis identified a combination of local composition-transition-distribution (LD) and conjoint triad (CT) descriptors among the five most informative features, as shown in Figure 7 and detailed in Table 13. The five highest-ranked features were indices 2451, 806, 1673, 1582, and 792. Feature 2451, the most important descriptor, represented the composition of the M/S/T/Y amino-acid group in region I (last 75%) of Protein 2, indicating a strong contribution from the regional abundance of these residues. Feature 806 corresponded to the M/S/T/Y-M/S/T/Y-M/S/T/Y conjoint triad pattern in Protein 2, highlighting the importance of specific local three-residue sequence patterns. The third-ranked feature, index 1673, represented a K/R → K/R transition in region J (middle 75%) of Protein 1, whereas feature 1582 represented the same K/R → K/R transition in region I (last 75%) of Protein 1. Feature 792 corresponded to the M/S/T/Y-H/N/Q/W-M/S/T/Y conjoint triad pattern in Protein 2. Collectively, these top-ranked features demonstrate that XGBoost primarily utilized regional amino-acid composition and transition patterns together with specific three-residue amino-acid group patterns for PPI prediction.

**Table 13.** Top five XGBoost features identified by feature-importance analysis.

| Rank | Feature Index | Type | Feature Description |
| --- | --- | --- | --- |
| 1 | 2451 | LD | M/S/T/Y composition in region I (last 75%) of Protein 2 |
| 2 | 806 | CT | M/S/T/Y-M/S/T/Y-M/S/T/Y triad pattern in Protein 2 |
| 3 | 1673 | LD | K/R → K/R transition in region J (middle 75%) of Protein 1 |
| 4 | 1582 | LD | K/R → K/R transition in region I (last 75%) of Protein 1 |
| 5 | 792 | CT | M/S/T/Y-H/N/Q/W-M/S/T/Y triad pattern in Protein 2 |

**Figure 7.**
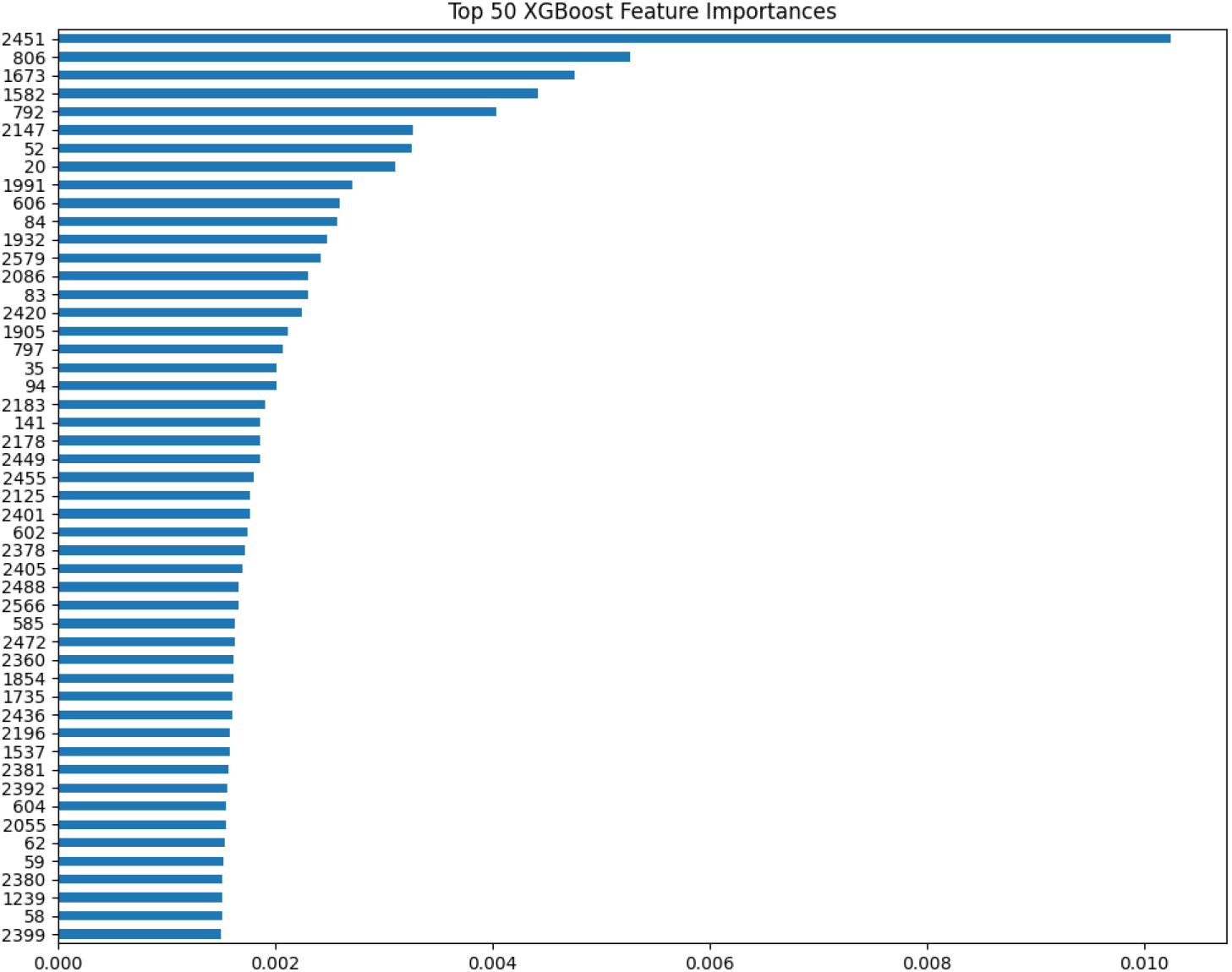
Top 50 XGBoost features ranked according to mean feature importance, highlighting Autocovariance (AC), Conjoint triad (CT), and Local Descriptors (LD) contributing to PPI prediction.

### 3.4 Discussion

The findings of this study demonstrate that coupling domain-informed, multi-scale sequence representations with classical ensemble learning architectures provides a robust, computationally efficient, and highly interpretable paradigm for protein-protein interaction (PPI) prediction. Although modern deep-learning models (e.g., DeepPPI^24^) can capture latent interaction representations, their black-box nature, high computational overhead, and susceptibility to overfitting on sparse interactome data present key limitations for translational applications. In contrast, our engineered 2,626 dimensional feature space, spanning 120 auto-covariance (AC) descriptors, 686 conjoint triad (CT) features (indices 121-806), and 1,820 local composition-transition-distribution (LD) features (indices 807-2626), which directly maps primary sequence topology to fundamental physicochemical properties governing molecular recognition.

A crucial finding of this work lies in the performance divergence observed between cross-validation and independent test-set evaluation. While XGBoost marginally surpassed Random Forest (RF) during five-fold cross-validation (97.97% versus 97.75% accuracy; F1-scores of 98.09% and 97.75%, respectively), RF demonstrated superior generalization on the unseen hold-out set, achieving peak performance across all metrics: 99.51% accuracy, 99.72% precision, 99.29% recall, 99.50% F1-score, an AUROC of 0.9990, and an AUPRC of 0.9989 (compared to XGBoost’s 98.54% accuracy, AUROC of 0.9979, and AUPRC of 0.9974). This highlights the importance of evaluating models beyond single-split partitions, as gradient boosting decision trees can over-index on subtle noise within training folds, whereas RF’s bootstrap aggregating (bagging) mechanism reduces variance and improves out-of-sample generalization across high-dimensional biological feature spaces.

Feature-importance attribution further elucidates the biochemical drivers of model predictions. Across both RF and XGBoost architectures, top-ranked features spanned three main functional categories: (i) short-range physicochemical correlations, where RF feature importance identified feature 52 (polarity *P*_1_ autocovariance of Protein 2 at lag 2) and feature 12 (hydrophobicity *H* autocovariance of Protein 2 at lag 2) as the top two most informative descriptors, indicating that local, spatial periodicity in polarity and hydrophobicity along the primary polypeptide backbone plays a key role in stabilizing protein-protein interfaces via electrostatic and hydrophobic interactions; (ii) local triad motif frequencies, where conjoint triad features such as feature 512 (A/G/V-M/S/T/Y-M/S/T/Y in Protein 2) and feature 806 (M/S/T/Y-M/S/T/Y-M/S/T/Y triad in Protein 2) were highly ranked by RF and XGBoost, respectively, reflecting the overrepresentation of flexible hydrophobic cores and hydrogen-bonding networks at interfacial contact surfaces; and (iii) regional amino-acid composition and transitions, where XGBoost feature attribution was strongly driven by local descriptor features, such as feature 2451 (M/S/T/Y composition in region I [C-terminal 75%] of Protein 2) and features 1673/1582 (K/R → K/R transitions in regions J and I of Protein 1), highlighting the functional role of basic, charged patches in establishing long-range electrostatic steering during complex formation.

Despite these strong predictive metrics, certain limitations of the present study warrant consideration: (i) while the gold-standard sequence-based PPI and Human-PPI datasets provided structured positive and negative interaction pairs, the absolute sample size remains modest relative to the vast, unannotated human interactome, meaning performance estimates may be influenced by dataset-specific negative sampling biases that can artificially inflate classification metrics compared to challenging proteome-wide screening scenarios; and (ii) sequence-based descriptors inherently collapse three-dimensional spatial arrangements into linear sequence profiles, so subtle conformational changes, post-translational modifications (PTMs), and transient quaternary dynamics are not explicitly modeled. Future studies should aim to address these limitations by incorporating structural representations from AlphaFold predictions^25^, evolutionary profile alignments (PSSMs), and expanding benchmark evaluations across larger, imbalance-corrected interactome datasets.

## 4 Conclusion

In conclusion, this study presents an end-to-end computational framework for protein-protein interaction (PPI) prediction that integrates multi-scale, biologically interpretable sequence descriptors with classical machine-learning classifiers. By combining 120 auto-covariance (AC), 686 conjoint triad (CT), and 1,820 local descriptors (LD) features, the pipeline effectively encodes local physicochemical correlations, triad motif distributions, and regional amino-acid transitions across interacting protein pairs. Rigorous evaluation across five-fold cross-validation and an independent unseen hold-out dataset established the performance and robustness of ensemble-based methods over traditional margin-based classifiers like Support Vector Machines (SVM, 87.58% test accuracy). Although XGBoost achieved superior cross-validation performance (97.97% accuracy), Random Forest demonstrated outstanding generalization on unseen test data, attaining 99.51% accuracy, 99.72% precision, 99.29% recall, a 99.50% F1-score, an AUROC of 0.9990, and an AUPRC of 0.9989. Feature attribution analysis systematically identified short-lag polarity/hydrophobicity autocovariances, interfacial triad motifs (A/G/V–M/S/T/Y–M/S/T/Y), and basic residue transitions (K/R → K/R) as primary drivers of PPI discrimination, providing clear biological transparency alongside high classification accuracy. While the model’s reliance on linear sequence representations and a bounded benchmark dataset highlights the need for future validation on larger, structural, and evolutionary multi-modal datasets, the framework’s low computational footprint, lack of GPU dependency during inference, and high biological interpretability make it well-suited for high-throughput interactome screening, target prioritization, and early-stage computational drug discovery workflows.

## Data Availability

The dataset used in this study is available at:

Training data set (Intra-0): https://doi.org/10.6084/m9.figshare.21591618.v4

Unseen Human-PPI dataset: https://www.kaggle.com/datasets/spandansureja/ppi-dataset

## Acknowledgements

The authors would like to express their sincere gratitude to all individuals who contributed their time, expertise, and support during the course of this research.

